# Fast voltage sensitivity in retinal pigment epithelium: sodium channels and their novel role in phagocytosis

**DOI:** 10.1101/223719

**Authors:** Julia K. Johansson, Teemu O. Ihalainen, Heli Skottman, Soile Nymark

## Abstract

Despite the discoveries of voltage-gated sodium channels (Na_v_) from a number of non-excitable cell types, the presence of Na_v_-mediated currents in cells of the retinal pigment epithelium (RPE) has been dismissed as a cell culture artifact. Here, we challenge this notion by demonstrating functional Na_v_1.4-Na_v_1.6 and Na_v_1.8 channels in human embryonic stem cell derived and mouse RPE. Importantly, we show that Na_v_s are involved in photoreceptor outer segment phagocytosis: blocking their activity significantly reduces the efficiency of this process. Consistent with this role, Na_v_1.8 co-localizes with the endosomal marker Rab7 and, during phagocytosis, with opsin. Na_v_1.4 localizes strongly to the cell-cell junctions together with the gap junction protein Connexin 43. During phagocytosis, both are localized to the phagosomes with a concurrent decrease in the junctional localization. Our study demonstrates that Na_v_s give the capacity of fast electrical signaling to RPE and that Na_v_s play a novel role in photoreceptor outer segment phagocytosis.

## Introduction

In the vertebrate eye, the retinal pigment epithelium (RPE) forms a barrier between the retina and the choroid^1–3^. Its cells are associated closely with photoreceptors: their apical sides surround the outer segments with long microvilli, and the basolateral sides are attached to Bruch’s membrane, an extracellular matrix separating the RPE from the choroid^3,4^. The RPE has many functions that are vital to retinal maintenance and vision, such as maintaining the visual cycle, secreting important growth factors, delivering nutrients to the photoreceptors from the blood stream while removing metabolic end products, and absorbing scattered light^1,3^. Most importantly, though, the RPE maintains ionic homeostasis in the subretinal space^5^ and sustains photoreceptor renewal by phagocytosing their shed outer segments^1,6^. Phagocytosis is highly essential for vision and it is under strict diurnal control, initiated at light onset for rods and at light offset for cones^7,8^. This evolutionary conserved molecular pathway is receptor mediated and precisely regulated, however the exact signaling cascades are still not completely understood^9^. Recent studies imply the importance of specific ion channels in this process including the L-type calcium currents linked to calcium-dependent potassium and chloride channels^10^.

Since the first single-cell recordings from RPE cells in 1988^11^, a large variety of different ion channels have been identified in them^5^. Much of this work was done on RPE cultured from various species, and some channels have been found only in cultured RPE^12–14^. Thus, there appear to be important differences between freshly isolated and cultured RPE^15^. One of the channel types identified in cultured RPE is the voltage-gated Na^+^ channel (Na_v_)^16,17^; Na_v_-mediated currents have not been recorded from freshly-isolated RPE^18,19^. Thus, it is widely thought that the expression of Na_v_ in RPE cells is the result of the neuroepithelial differentiation in culture^5,15,20^.

Here, we challenge this conventional wisdom by demonstrating functional Na_v_ channels in cultured human embryonic stem cell (hESC) -derived RPE *in vitro* and freshly-isolated mouse RPE *in vivo.* Our work reveals that Na_v_ channels in RPE are primarily localized to the tight junctions that connect the cells. We conclude that this specific localization of Na_v_s to cell-cell contacts accounts for previous findings that dissociated RPE cells lack Na_v_-mediated currents. Specifically, we hypothesize that Na_v_ channels could co-regulate photoreceptor outer segment (POS) phagocytosis. Our hypothesis is supported by a recent demonstration of the involvement of Na_v_ channels in phagocytosis of mycobacteria by macrophages^21^. Our work provides evidence that Na_v_1.8 localizes with the endosomal marker Rab7, and that they both accumulate with the phagosomal particles. Further, Na_v_1.4 co-localizes with connexin 43 (Cx43), a constituent of the gap junctions connecting RPE cells^22^. During phagocytosis, however, they both are localized to the phagosomes with a concomitant decrease in junctional localization. Interestingly, this phagosomal translocation was significantly reduced by selective Na_v_ channel blockers. Moreover, the selective blockers combined with the universal Na_v_ blocker tetrodotoxin (TTX) reduced the total number of POS particles by up to 34%. More generally, our observations add to the growing body of evidence that Na_v_s play diverse roles in a variety of classically non-excitable cell types ranging from astrocytes and microglia to macrophages and cancer cells (for review, see^23^).

## Results

### Functional voltage-gated sodium channels are present in RPE monolayer cultures derived from human embryonic stem cells

We used whole-cell recordings from mature hESC-derived RPE in K^+^ free extra- and intracellular solutions to observe transient inward currents elicited by a series of depolarizing voltage pulses after strong hyperpolarization to -170 mV (Fig. 1c, n=33). These recordings were done from an intact monolayer (Fig. 1a) in the presence and absence of a gap-junction antagonist (18α-glycyrrhetinic acid). Similar currents were also identified, albeit infrequently and with reduced amplitudes, in cells from freshly dissociated mature hESC-derived RPE (Fig. 1b, 1d, n=6). In all experiments, the current resembled the Na_v_ current characteristic of excitable cells: it had the typical current – voltage relationship (Fig. 1e) and showed fast activation and inactivation (Fig. 1c). The current was activated at about -50 mV and peaked at about -13 mV with a maximum amplitude of 294 ± 79 pA (mean ± SEM, n=18). Taking into account the average membrane capacitance 39 ± 5 pF (mean ± SEM), the average current density was 8.7 ± 3.4 pA/pF (mean ± SEM, n=18). To investigate the time dependency of recovery from inactivation, we used a paired-pulse protocol (Fig. 1f). The current was recorded after a second depolarizing pulse given at increasing time intervals until it finally recovered to its full size (Fig. 1f). The second peak currents were subsequently normalized to the prepulse peak current and plotted against the time between the two voltage pulses (Fig. 1g). Our data was fitted with an exponential function and the best fit yielded to τ = 47 ms (n=3).

**Figure 1.**
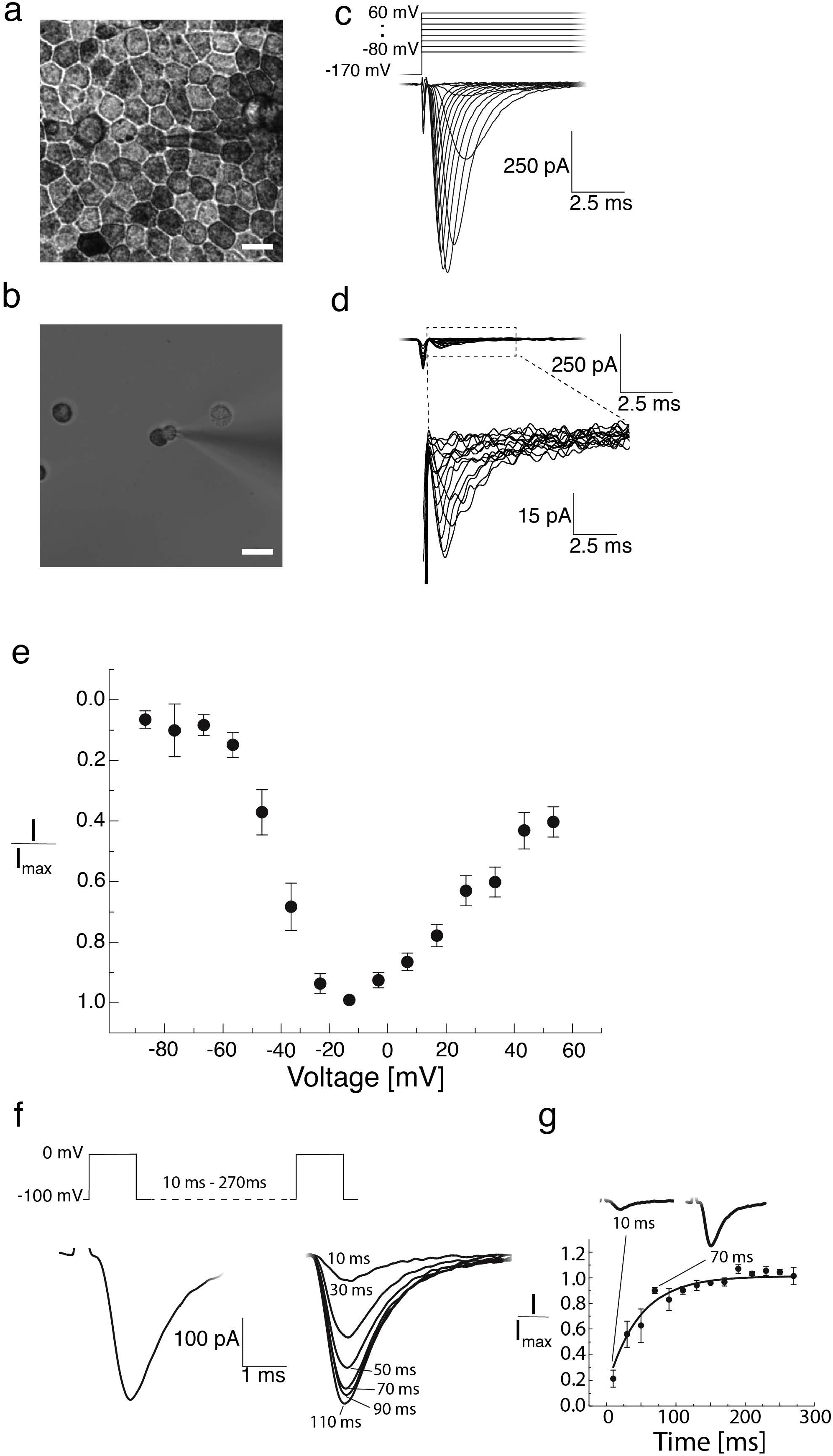
Patch clamp recordings of Na^+^ currents from hESC-derived RPE in different cell morphologies. Brightfield light microscopy images of hESC-derived RPE cells. (**a**) Mature hESC-derived RPE grown on insert for 2 months showing strongly pigmented cells and characteristic epithelial morphology. (**b**) Mature hESC-derived RPE was dissociated yielding single cells with typical morphology showing pigmented apical and non-pigmented basal sides. The cells were seeded on poly-L-lysine coated glass coverslip and let to adhere for 10 min before recording. Whole-cell patch clamp recordings as responses to a series of depolarizing voltage pulses (-80 to +60 mV) after strong hyperpolarization either (**c**) from mature monolayer of hESC-derived RPE or (**d**) from single hESC-derived RPE cells. Patch clamp pipette is visible in the middle of the images. Scale bars 10 μm. (**e-g**) Analysis of the monolayer recordings. (**e**) The average normalized peak current – voltage relationship (I/I_max_ vs V_m_, n=33). Data points indicate mean±SEM. (**f-g**) The time dependency of recovery from inactivation. The second peak currents were normalized and plotted against the voltage pulse interval (10–270 ms). The best fit to an exponential function was obtained with τ = 47 ms (n=3).

The presence of Na_v_ currents was confirmed using the universal extracellular Na_v_ channel blocker tetrodotoxin (TTX). Addition of 1 μM TTX to the bath reduced the amplitude of the current to roughly one half of that recorded in the control extracellular solution (Fig. 2a). Thus, the recorded current was sensitive to TTX but required reasonably high concentrations. Furthermore, the sensitivity to TTX varied between the cells and in some cases even 10 μM TTX was not enough to block the current (Fig. 2a). On the other hand, 2mM QX-314, an intracellular Na_v_ channel blocker added to the internal solution of the patch pipette, removed the current rapidly after breaking into the whole-cell configuration (Fig. 2a).

**Figure 2.**
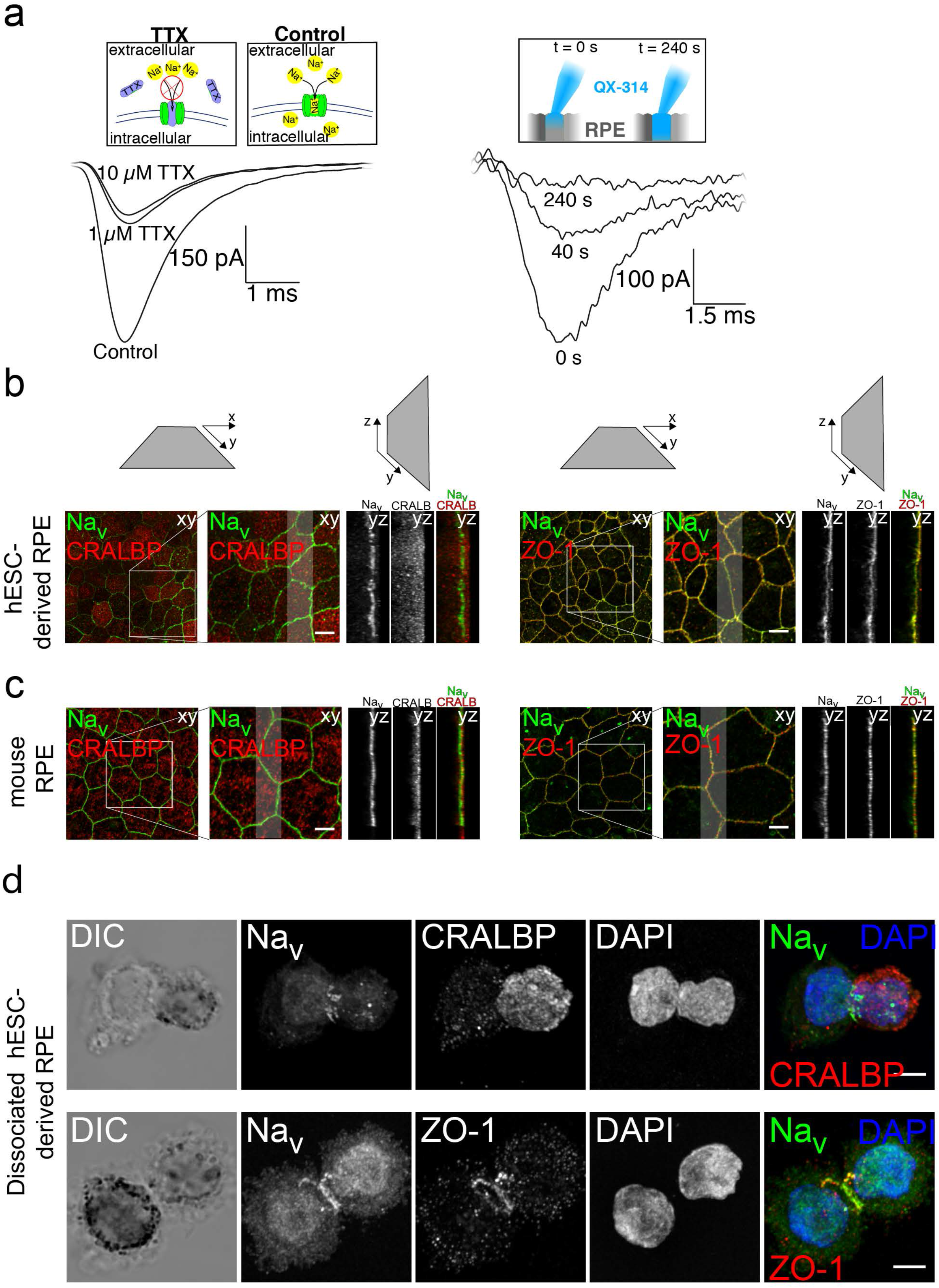
Blocker sensitivity and distribution of Na_v_ channels. Patch clamp recordings were performed on mature hESC-derived RPE monolayers. (**a**) Applying TTX extracellularly (either 10 μM or 1 μM) did not entirely block the current (left). The current was completely removed by intracellular QX-314 (2mM) (right). Laser scanning confocal microscopy (LSCM) images on Na_v_ distribution in RPE cells showing different imaging planes of the confocal data. LSCM data Z-maximum intensity projections (Z-MIP) of (**b**) hESC-derived and (**c**) mouse RPE stained against Na_v_ channels (green) and RPE marker CRALBP (red) (left) or tight junction marker ZO-1 (red) (right), together with cross-sectional X-MIPs from the highlighted regions. (**d**) Dissociated hESC-derived RPE cells were let to adhere to poly-l-lysine coated coverslips for 30 min, fixed and immunolabeled against Na_v_ together with RPE marker protein CRALBP (up) or tight junction marker ZO-1 (down). The Na_v_ label concentrated on the belt-like region in the middle of the cell, between the basal and apical sides. Scale bars 5 μm.

### Voltage-gated sodium channels localize to the apical side of the tight junctions in hESC-derived as well as mouse RPE

Our patch-clamp data indicated that functional Na_v_ channels are present in the hESC-derived RPE cells. We also wanted to verify the expression *in situ* and study the precise cellular localization of the channels by performing immunofluorescence studies: the cellular retinaldehyde binding protein (CRALBP), a marker for mature RPE cells^24,25^, was labelled together with the universal Na_v_ channel marker. These hESC-derived RPE samples were then imaged with a laser scanning confocal microscope (LSCM) by acquiring 3D image stacks (Fig. 2b) and the data were denoised by deconvolution. This showed that Na_v_ was present in fully differentiated RPE. Furthermore, the Na_v_ label concentrated primarily on the cellular borders while the CRALBP label was more uniformly localized to the apical side of the hESC-derived RPE (Fig. 2b).

Since the Na_v_ label seemed to localize to the cell-cell junctions, immunolabelling was next carried out with the tight junction marker ZO-1. The labels showed highly overlapping distributions, suggesting that Na_v_ localizes mostly into the tight junctions (Fig. 2b). The expression of Na_v_ channels in RPE has previously been thought to be induced *in vitro* by the cell culturing^18,19^, therefore we wanted to confirm the presence of the Na_v_ channels *in vivo* by using freshly isolated and non-cultured mouse RPE (Fig. 2c). The same antibodies gave highly similar labeling distributions in mouse RPE as in hESC-derived RPE: the CRALBP label was cytoplasmic on the apical side of the cells while ZO-1 and Na_v_ concentrated on the cell-cell junctions (Fig. 2c).

We investigated the mechanism underlying the disappearance of Na_v_ currents from acutely isolated RPE cells (Fig. 1d) and discovered evidence for internalization following destruction of the tight junctions. In dissociated hESC-derived RPE cells seeded for 30 min on glass and immunolabeled with the universal Na_v_ marker, CRALPB and ZO-1, the Na_v_ label was primarily concentrated in the narrow region separating the apical and basolateral sides of the cell. ZO-1 and Na_v_ formed a clear ring-like structure between the apical and basal membranes following relaxation of junctional tension, suggesting that cytoskeletal contractility is involved in the internalization process (Fig. 2d). Due to the junctional disruption, Na_v_s might not be accessible to pass ionic currents.

### RPE cells express voltage-gated sodium channel subtypes Na_v_1.4, Na_v_1.5, Na_v_1.6 and Na_v_1.8

Our previous experiments indicated that functional voltage-gated sodium channels are present in RPE. Since ten different Na_v_ channel subtypes have been identified, Na_v_1.1 – Na_v_1.9 and Nax, with drastically different expression profiles in diverse cell types, we wanted to investigate which specific channel subtypes are functionally expressed in the RPE cells. At the mRNA level, previous work has shown expression of several Na_v_ channels in donated human RPE, specifically Na_v_ subtypes 1.2–1.6 and Na_v_1.9^26,27^. We performed immunolabeling experiments with mouse and hESC-derived RPE using specific antibodies against channel subtypes Na_v_1.1 – Na_v_1.9 (Fig. 3a, 3b, Fig. S1). Confocal microscopy showed that Na_v_1.4 localizes as beads-on-a-string to the cell-cell junctions and strongly co-localizes with the gap junction marker connexin 43 (Cx43) (Fig. S3). Na_v_1.8, on the other hand, localized overall to the apical side of the RPE cells (Fig. 3a, 3b). These data indicated that especially the Na_v_1.4 and Na_v_1.8 channels which are usually expressed in skeletal muscle and dorsal root ganglia^28,29^, respectively, are also present in RPE cells. Moreover, subtypes Na_v_1.5 and Na_v_1.6, the predominant channels of cardiac muscle, and the adult central nervous system, respectively^30^, were also identified. However, the expression of Na_v_1.5 was punctate and not detectable in all cells whereas Na_v_1.6 showed a more homogenous labeling pattern and uniform expression between the cells (Fig. 3a, 3b). Subtypes 1.1–1.3, 1.7 and 1.9 were not detected (Fig. S1). Additionally, we investigated the changes in channel subtype localization patterns during maturation of hESC-derived RPE (Fig. S2). The immunolabeling experiments indicated that the subtypes Na_v_1.4, Na_v_1.5 and Na_v_1.8 changed from homogeneous cellular distribution to more specific localization either to cell-cell junctions (Na_v_1.4) or to the apical side of the epithelium (Na_v_1.5 and Na_v_1.8) during the first 9 days of maturation (Fig. S2).

**Figure 3.**
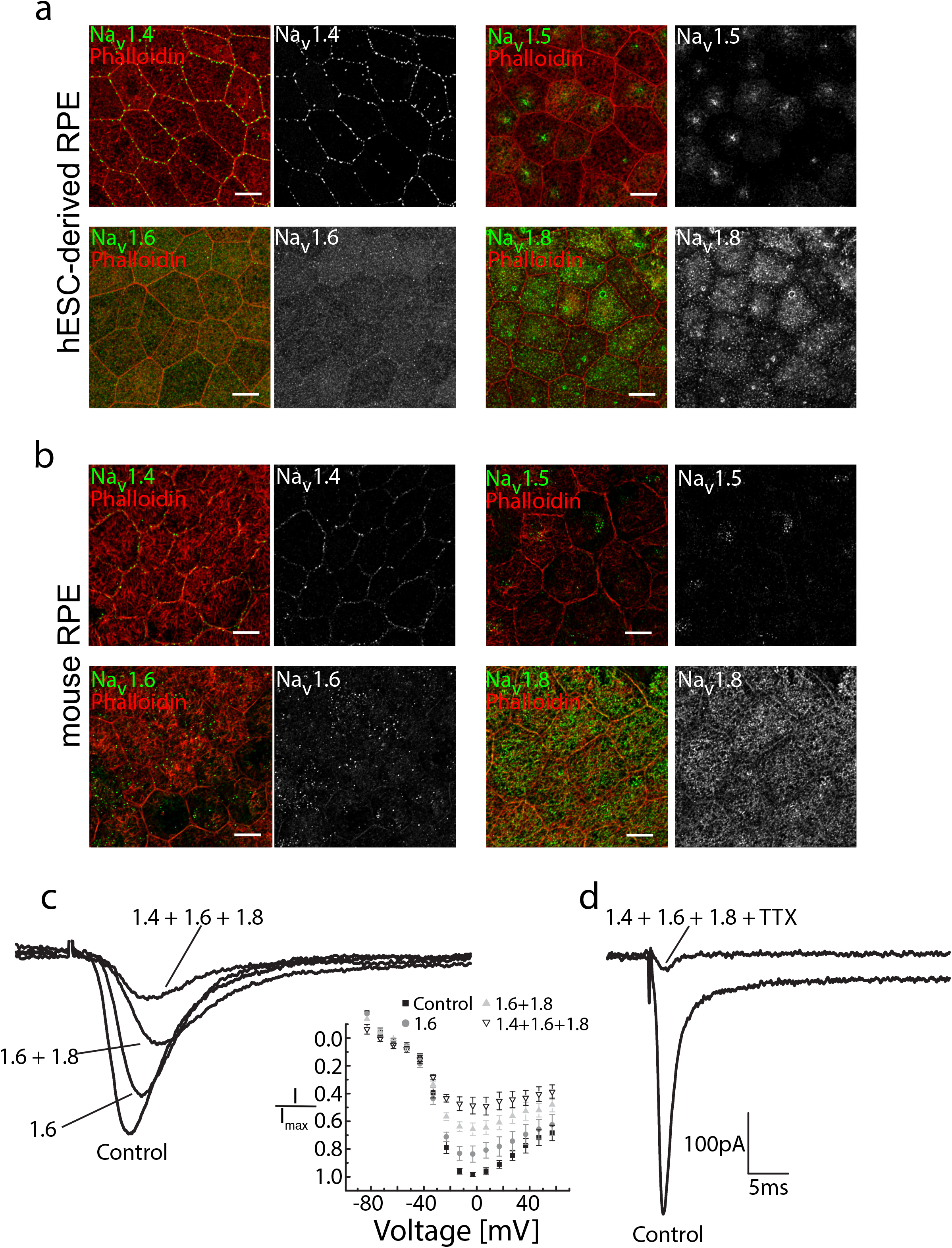
Immunolabeling of different Na_v_ subtypes in hESC- and mouse derived RPE, and patch clamp recordings with selective Na_v_ blockers. The specific pattern of Na_v_ subtypes was studied by immunolabeling. (**a**) Laser scanning confocal microscopy Z-maximum intensity projections of mature hESC-derived or (**b**) mouse RPE. Na_v_ subtypes 1.4–1.6 and 1.8 (green) were immunolabeled together with filamentous actin (phalloidin stain, red). Patch clamp recordings were performed on mature hESC-derived RPE using selective blockers for channel subtypes. (**c**) Na_v_ subtypes were sequentially blocked by extracellularly applied 4,9-AnhydroTTX (30 nM, Na_v_1.6 blocker) in combination with A-803467 (1 μM, Na_v_1.8 blocker) and μ-Conotoxin GIIB (600 nM, Na_v_1.4 blocker). The average normalized peak current – voltage relationship (I/I_max_ vs V_m_) was determined from all recordings (n=7). (**d**) Applying the selective blockers in combination with TTX (10 μM) removed most of the Na_v_ current. Scale bars 10 μm.

To verify further the functional expression of the most prominent channel subtypes by electrophysiology, we repeated our patch clamp recordings using highly selective blockers for the channels Na_v_1.4 and Na_v_1.8 and a less specific blocker for Na_v_1.6. At the time of our study, to our knowledge there was no selective blocker available for Na_v_1.5 without potential cross-reactivity. The average current-voltage relationship (I-V curve) was determined from all these recordings (n=7) (Fig. 3c). The current was sensitive to the combination of 30 nM 4,9-anhydro-TTX (Na_v_1.6 blocker), 1 μM A-803467 (Na_v_1.8 blocker) and 600 nM μ-conotoxin GIIB (Na_v_1.4 blocker) (Fig. 3c) and the effect of inhibition was more potent with each blocker thus confirming the expression and functionality of these channel subtypes in the hESC-derived RPE. However, the effect of inhibition was more significant when the blockers were combined with 10μM TTX (Fig. 3d).

### Inhibition of voltage-gated sodium channels significantly reduces the number of POS particles in hESC-derived RPE

Our previous experiments showed that Na_v_1.4, Na_v_1.5, Na_v_1.6 and Na_v_1.8 are expressed both in mouse and mature hESC-derived RPE. However, their physiological relevance remained unknown. Phagocytosis of POS is one of the major roles of RPE^3^, and a plausible candidate function for the Na_v_ channels, as it requires rapid activation and high synchronization^31^. Thus, next we wanted to investigate the potential importance of Na_v_ channels for POS phagocytosis *in vitro*. Our phagocytosis assays and subsequent labeling with opsin and phalloidin indicated that the hESC-derived RPE cells have the ability to bind and take up purified POS particles isolated from porcine eyes (Fig. 4a). To study the involvement of Na_v_ channels in this pathway, the assay was repeated in the presence of the highly selective Na_v_ 1.4 and 1.8 blockers and TTX. Interestingly, the immunolabeling with opsin and ZO-1, a tight junction protein and cell border marker, showed a reduction in the total number of bound and internalized POS particles (Fig. 4b). To quantify the effect, large fields were imaged from each blocker condition and the number of POS particles was compared with controls. The results showed that the selective blockers caused a 13% (Na_v_ 1.4 blocker, n= 10), 27% (Na_v_ 1.8 blocker, n=14) or 34% (Na_v_ 1.4 and Na_v_ 1.8 blockers together with TTX, n=18) reduction in the total number of POS particles labeled with opsin (Fig. 4c).

**Figure 4.**
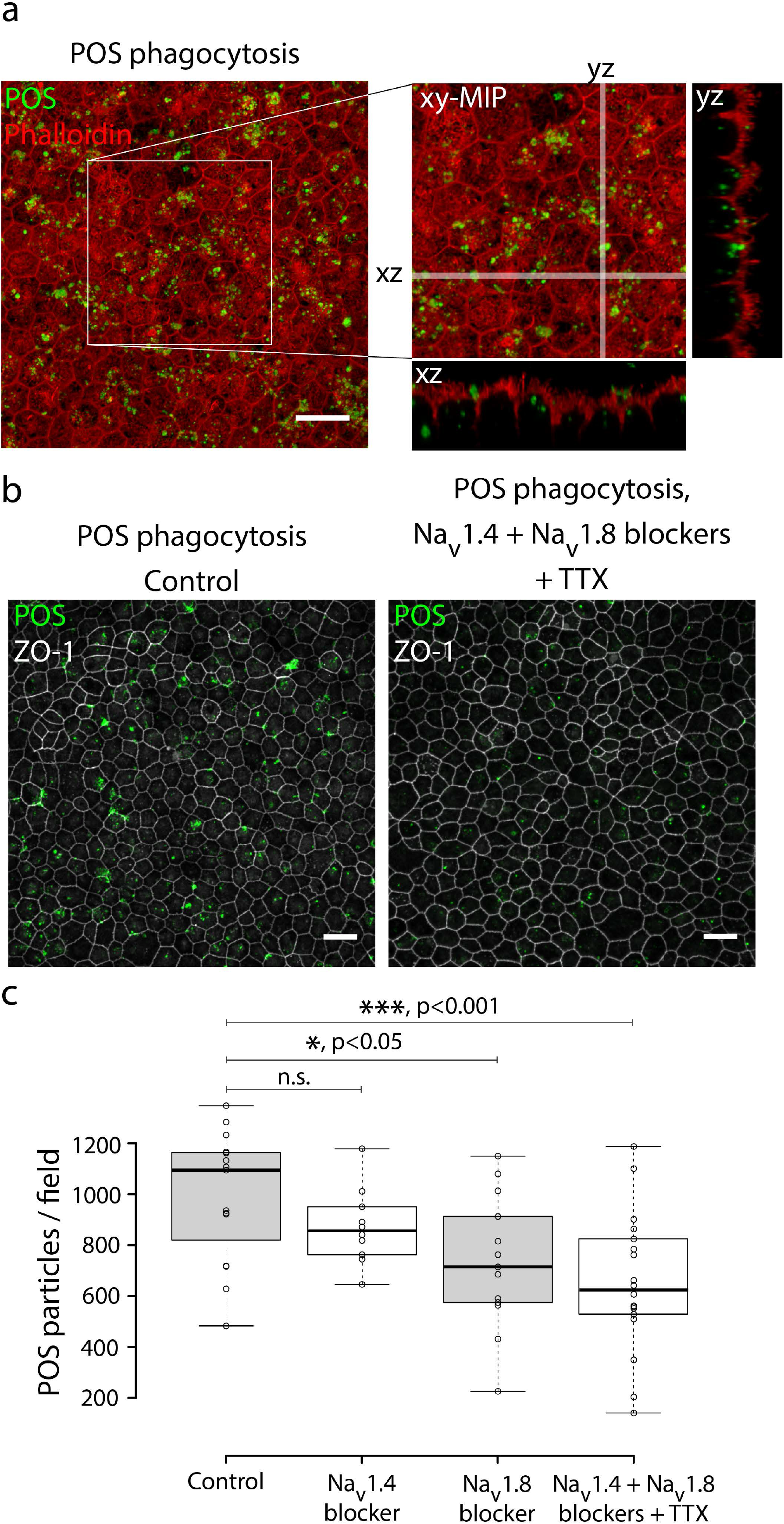
POS phagocytosis assay of hESC-derived RPE with selective Na_v_ blockers. POS phagocytosis assay was performed on mature hESC-derived RPE by incubating the monolayers with purified porcine POS particles. (**a**) Laser scanning confocal microscopy (LSCM) Z-maximum intensity projections (Z-MIP) of the POS particles (green) with filamentous actin staining (red) after 2h of phagocytic challenge. (**b**) LSCM Z-MIP images of ZO-1 (grey) together with opsin (green). The assay with selective Na_v_ blocker combination reduced the total number of POS particles. Scale bars 20 μm. (**c**) Quantification of control (n=15) and blocker samples showed a 13% (Na_v_1.4 blocker, n= 10), 27% (Na_v_1.8 blocker, n=14) or 34% (Na_v_1.4 and Na_v_1.8 blockers with TTX, n=18) reduction in the number of POS particles labeled with opsin. Center lines show the medians; box limits indicate the 25th and 75th percentiles as determined by R software; whiskers extend to minimum and maximum values.

### Voltage-gated sodium channels Na_v_1.4 and Na_v_1.8 are involved in POS phagocytosis in mouse RPE *in vivo*

The experiments with pharmacological inhibitors of Na_v_ channel subtypes Na_v_1.4, and Na_v_1.8 indicated that these channels are involved in POS particle uptake. To investigate further their role in the phagocytosis process *in vivo*, we performed immunolabeling experiments with mouse eyes that had been prepared either at light onset near the diurnal peak of phagocytosis or 10 h after light onset. The role of the channels in POS uptake was studied by comparing the immunolabeling of Na_v_1.4, Na_v_1.8 and opsin. At light onset both channels localized to the bound POS particles (Fig. 5a). Since previous experiments had indicated that Na_v_1.4 localized strongly to gap junctions (Fig. S3c), we labeled it together with Cx43 as well as opsin. Co-localization of these three markers was most evident at light onset (Fig. 5b), yet we observed a reduction in Na_v_1.4 and opsin co-localization at 2 h time point (Fig. 5c). Interestingly, after 10 h, when phagocytosis process is over^32^, both Na_v_1.4 and Cx43 stains were again detected at cell-cell junctions (Fig. 5d).

**Figure 5.**
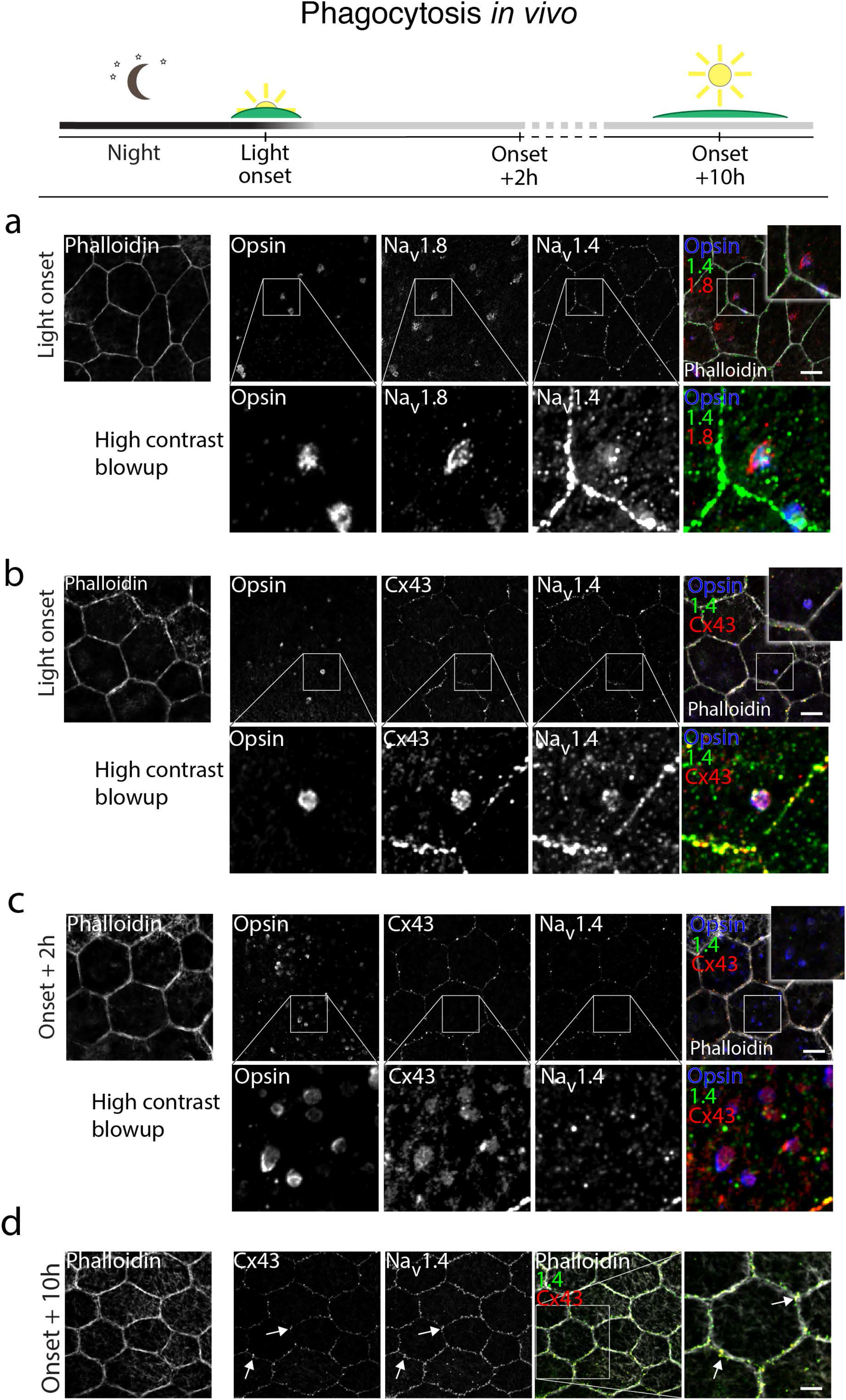
POS phagocytosis *in vivo* and the roles of Na_v_1.4 and Cx43. Phagocytosis was studied *in vivo* by dissecting mouse eyes at various time points during the circadian cycle. Filamentous actin was stained with phalloidin (gray in the merged image) to highlight epithelial cell-cell junctions and lower panels show a high contrast blow up of selected regions. Laser scanning confocal microscopy Z-maximum intensity projections of mouse RPE prepared at light onset showed a localization of opsin labeled POS particles (blue) and Na_v_1.4 (green) together with (**a**) Na_v_1.8 (red) or (**b**) Cx43 (red). (**c**) 2 h after the light onset, labeling of both Na_v_1.4 (green) and Cx43 (red) in the cell-cell junctions was dramatically reduced and Cx43 localized with POS particles. (**d**) 10 h after the light onset, Na_v_1.4 (green) and Cx43 (red) both localize to cell-cell junctions. Scale bars 10 μm.

We next wanted to investigate further the role of Na_v_1.8 in the POS phagocytosis. For this purpose, we combined the labeling of Na_v_1.8 with Rab7, a GTPase which is generally recognized in the regulation of vesicle docking and fusion, specifically with late endosomes and lysosomes^33,34^. In macrophages Na_v_1.5 has previously been demonstrated to preferentially localize with Rab7 rather than the endosomal markers early endosome antigen 1 (EEA1) or lysosomal-associated membrane protein 1 (LAMP-1)^35^. To date, controversy persists as to whether Rab7 is additionally involved in the earlier steps of endosomal trafficking^36^. We investigated how Na_v_1.8 labeling correlated with Rab7 and EEA1 as well as the phagocytosis markers MerTK and integrin αvβ5 (Fig. S4, Fig. 6). Na_v_1.8 was found to partially co-localize with EEA1 and integrin αvβ5, however the distribution similarity was most evident with Rab7 (Fig. S4, Fig 6). Moreover, both Na_v_1.8 and Rab7 also localized with opsin throughout the 2 h follow-up of phagocytosis (Fig. 6a, 6b, Fig. 5a). When the peak of phagocytosis was over (10 h time point), the overall labelling pattern of Na_v_1.8 appeared significantly more homogeneous (Fig. 6c).

**Figure 6.**
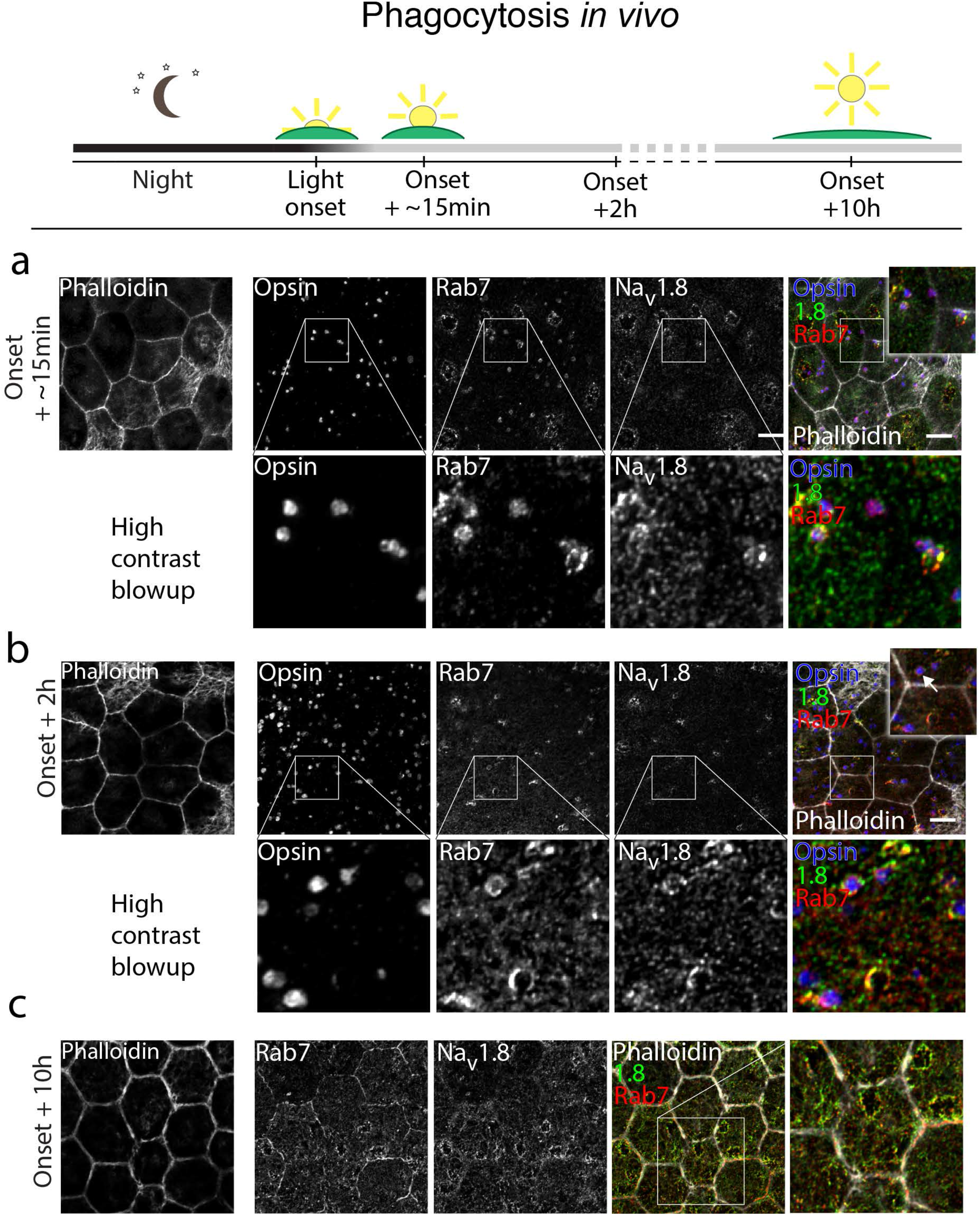
POS phagocytosis *in vivo* and the roles of Na_v_1.8 and Rab7. The role of Na_v_1.8 and its interaction with endosomal marker Rab7 was studied by dissecting mouse eyes at various time points during the circadian cycle. Filamentous actin was stained with phalloidin (gray in the merged image) to highlight epithelial cell-cell junctions and lower panels show a high contrast blow up of selected regions. Laser scanning confocal microscopy Z-maximum intensity projections of mouse RPE prepared (**a**) at 15 min after the light onset indicated clustering of Rab7 (red) and Na_v_1.8 (green) together with opsin (blue) labeled POS particles. (**b**) 2 h after the light onset, Na_v_1.8 (green) and Rab7 (red) still co-localize with POS particles but start to show more homogeneous localization pattern. (**c**) 10 h after light onset a diffuse labeling was shown for both and Rab7 (red) and Na_v_ 1.8 (green). Scale bars 10 μm.

The redistribution of Na_v_ channels occurring during phagocytosis (Fig. 7a) was studied *ex vivo* with the channel blockers (Fig. 7b). For this purpose, we developed an assay where freshly opened mouse eyecups were incubated in physiological conditions with blocker solutions for 1 h starting at 15 min prior to light onset. The blocker for Na_v_1.4 as well as the combination of all Na_v_ blockers significantly prevented the redistribution of Na_v_1.4 compared to the control (Fig. 7b). The inhibition effect was similarly observed in hESC-derived RPE *in vitro* when the POS challenge was carried out with blocker solutions (Fig. 7c). We did not observe significant differences in the labeling pattern of Na_v_1.8 or Rab7 after the blocker incubation. Taken together, these experiments demonstrate the participation of Na_v_ channels in the phagocytotic processes of RPE cells *in vitro* and *in vivo.*

**Figure 7.**
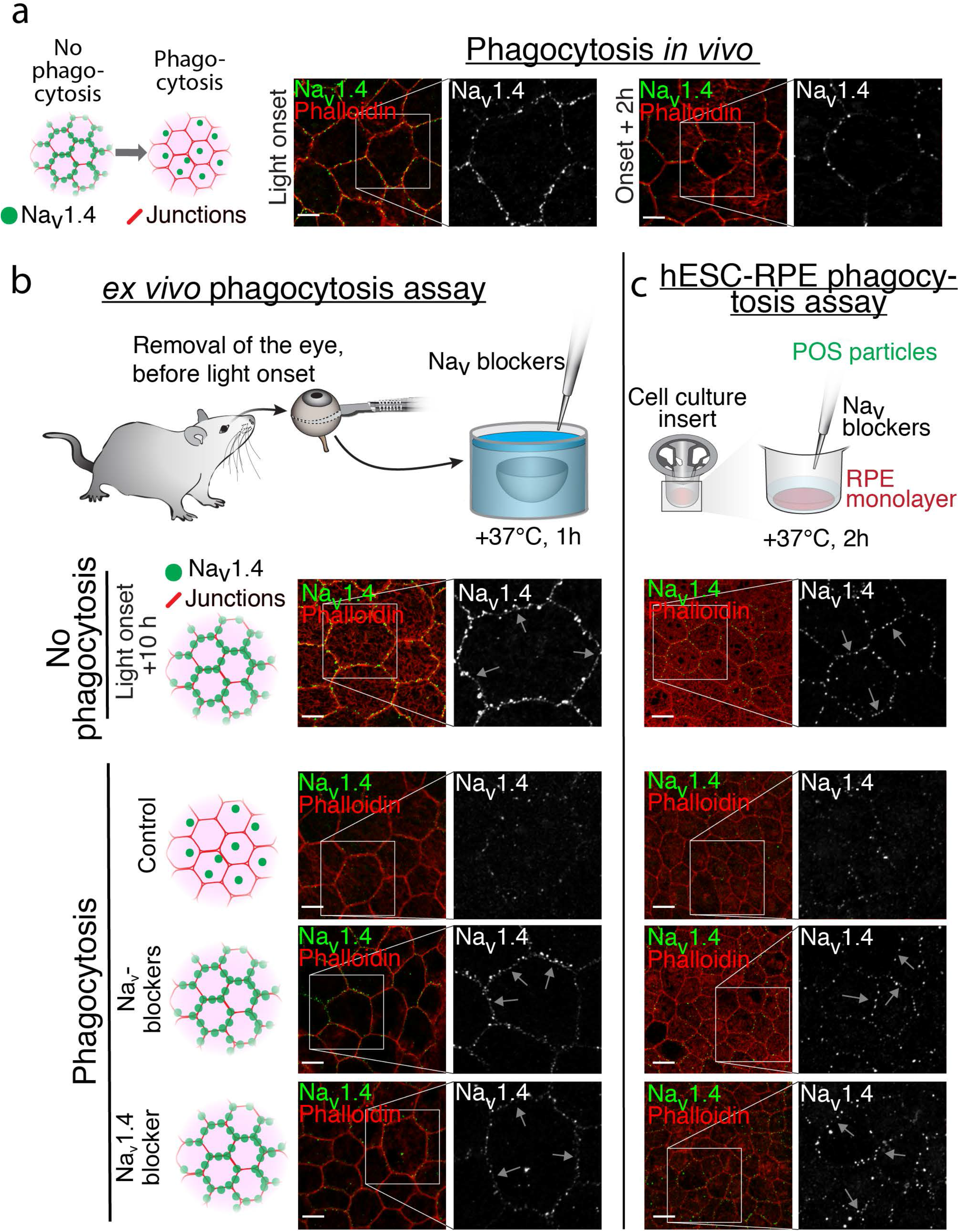
Redistribution of Na_v_1.4 during POS phagocytosis. The redistribution of Na_v_1.4 during phagocytosis and the effect of Na_v_ blockers was studied in mouse and hESC-derived RPE. Filamentous actin was stained with phalloidin (red) to highlight epithelial cell-cell junctions. Laser scanning confocal microscopy Z-maximum intensity projections of (**a**) Na_v_1.4 localization at light onset and 2 h after showed disappearance of the beads-on-a-string type labeling from cell-cell junctions. Different assays were used to investigate Na_v_1.4 distribution during phagocytosis and the effect of selective blockers for Na_v_1.4 and Na_v_1.8 in combination with TTX (Na_v_ blockers), or only of the selective blocker for Na_v_1.4. (**b**) The redistribution of Na_v_1.4 was studied *ex vivo* by incubating opened eyecups in control solution or with the selective blockers. In both of the blocker samples, the redistribution was inhibited and the beads-on-a-string type labeling remained visible (highlighted by arrows) in the cell-cell junctions. (**c**) The hESC-derived RPE phagocytosis assay showed a highly similar redistribution of Na_v_1.4 and the blockers had the same effect *in vitro.* Scale bars 10 μm.

## Discussion

Recent studies in diverse cell types have revolutionized our understanding of the roles that Na_v_ channels have in cellular functions; no longer are these proteins considered important only in “classically” electrically excitable tissues. Here, we provide the first evidence that Na_v_ channels are localized to the tight junctions between the epithelial cells (Fig. 2) and that their activity co-regulates phagocytosis in RPE. Our observations of Na_v_ channels and Na_v_-mediated currents in intact RPE preparations (mature hESC-derived RPE monolayers and freshly isolated mouse RPE) demonstrate that previous observations of Na_v_-mediated currents in cultured RPE cells are not preparation-dependent artifacts^19^. Rather, the absence of Na_v_-mediated currents in acutely isolated RPE cells (Fig. 1d) likely results from the destruction of tight junction complexes during dissociation (Fig. 2d). Internalization of Na_v_ channels, of course, would result in diminution or absence of membrane currents mediated by these channels as observed by us (Fig. 1d) and others^18^. The observation of Na_v_ currents in recordings from hESC-derived RPE monolayers, we believe, is strong evidence that cells in RPE with intact tight junctions usually express functional Na_v_s in their plasma membranes. Unfortunately, the denser and longer apical microvilli in mouse RPE prevented us from performing recordings from intact mouse tissue.

The properties of Na_v_-mediated currents in hESC-derived RPE cells are consistent with observations in other non-neuronal cells. Specifically, the relative insensitivity of the channels to TTX (block by 10 μM TTX) is similar to that of Na_v_s in macrophages^37^ and in cultured cancer cells^38^. Additionally, it is consistent with our observation that RPE cells were labeled strongly by an anti-Na_v_1.8 antibody; Na_v_1.8 is the least sensitive of Na_v_ channels to TTX^39^. Our pharmacological analysis of Na_v_-mediated currents using Na_v_ subtype-specific blockers and immunohistochemical analysis (Fig. 3) provided additional evidence for Na_v_1.8 expression along with lesser expression of Na_v_1.5.

A role for Na_v_ channels in POS phagocytosis is consistent with previous studies showing that Na_v_1.5 and Na_v_1.6 channels enhanced mycobacteria processing in human macrophages and that TTX attenuated phagocytosis in microglia^21,37,40,41^. The mechanisms by which voltage-gated channels with rapid activation and inactivation kinetics regulate phagocytosis are unclear. Although inhibition of Na_v_ channels did not abolish phagocytosis here—the observed 34% attenuation (Fig. 4c) was less dramatic than the previously reported effect of TTX in microglia^42^—Na_v_ channel activity clearly modulated phagocytosis as indicated by observations of bound POS particles. Furthermore, the channels could be involved directly in the circadian control of the pathway, as has been recently shown for other ion channels^10^.

Our co-labeling studies give a strong indication that Na_v_1.4 consistently interacts with Cx43 (Fig. 5). Similarly, an interaction between Na_v_ channels and Cx43 has been suggested for Na_v_1.5 in cardiac myocytes (for review, see^43^). Both Na_v_1.4 and Cx43 have previously been implicated in phagocytosis in macrophages, although the findings have been controversial^44,45^. We observed a redistribution of Na_v_1.4 and Cx43 from tight junctions during phagocytosis and for Cx43 the localization with opsin was still evident 2 h after light onset, near the peak expression level of a phagocyte cell surface receptor tyrosine-phosphorylated MerTK^32^. The involvement of Na_v_1.4 was supported by the fact that following Na_v_ blocker incubation, we observed a decrease in its translocation (Fig. 7) with a concurrent reduction in the number of POS particles (Fig. 4c). As with Na_v_1.4 and Cx43, Na_v_1.8 and Rab7 showed consistent co-localization (Fig. 6), and their cellular localization varied throughout phagocytosis during which both also localized with opsin. It is possible that the role of Na_v_1.8 and Rab7 is related to the regulation of endosomal processing or acidification of the endosomal pH, as has been suggested for Na_v_1.5 of macrophages in multiple sclerosis lesions^35^.

The fact that RPE expresses such a versatile array of Na_v_ channels suggests that besides phagocytosis, these channels also have other roles in the physiology of the RPE. Epithelial cells, including RPE, show strong calcium waves in response to mechanical stimulation^46–48^, and it is possible that Na_v_ channels serve to amplify or accelerate voltage changes. Moreover, it is well established that the Ca^2+^ binding protein calmodulin (Cam) interacts directly with the C-terminal domain of Na_v_^49^, and it was recently shown that the Ca^2+^ -free form of Cam, ApoCam, enhances the Na_v_ channel opening by several-fold^50^. Thus, Na^+^- and Ca^2+^-dependent signaling pathways can interact in epithelia as has been reported in the case of astrocytes^51^

In this study, we have demonstrated that Na_v_ channels can be identified in both hESC-derived and mature mouse RPE. This strongly indicates that expression of these channels is not associated with neuroepithelial differentiation nor due to specific culturing conditions. The channels are in fact implicated in POS phagocytosis, one of the key functions of RPE, which is essential for the survival of the retina. Further studies are required to elucidate the additional roles these channels have in the tissue, but it is evident that they are a vital part of the physiology of RPE and may thus contribute to the development of retinal diseases.

## Methods

### Cell culturing

Human ESC lines Regea08/023 and Regea08/017 were cultured as previously described^52^. Briefly, the hESC-derived RPE were spontaneously differentiated in floating cell clusters. The pigmented areas were isolated manually and the cells were dissociated with Tryple Select (1X, ThermoFisher Scientific) and filtered through cell strainer (BD Biosciences, NJ, USA). The isolated cells were then seeded on collagen IV-coated (human placenta, 5 μg/cm^2^; Sigma-Aldrich, MO, USA) 24-well plates (NUNC, Thermo Fisher Scientific, Tokyo, Japan) for enrichment. Subsequently, the pigmented cells were replated on collagen IV-coated (5 μg/cm^2^) culture inserts (Millicell Hanging Cell Culture Insert, polyethylene terephthalate, 1.0 μm pore size, EMD Millipore, MA, USA) for maturation. The cells were cultured at 37°C in 5 *%* CO^2^ in culture medium consisting of: Knock-Out Dulbecco’s modified Eagle’s medium (KO-DMEM), 15 % Knock-Out serum replacement (KO-SR), 2 mM GlutaMax, 0.1 mM 2-mercaptoethanol (all from Life Technologies, Carlsbad, CA), 1 % Minimum Essential Medium nonessential amino acids, and 50 U/mL penicillin/streptomycin (from Cambrex BioScience, Walkersville,MD). The culture medium was replenished three times a week. Mature monolayers typically showed transepithelial resistance values (TER) of over 200 Ω cm^2^.

The National Authority for Medicolegal Affairs Finland approved the study with human embryos (Dnro 1426/32/300/05). The supportive statement from the local ethics committee of the Pirkanmaa hospital district Finland allows us to derive and expand hESC-lines from surplus embryos excluded from infertility treatments and to use the lines for research purposes (R05116). Novel cell lines were not derived in this study

### Sample preparation

For monolayer patch clamp recordings and immunolabeling, the membrane of the culture insert was removed from the insert holder and cut into smaller pieces. The cells were rinsed three times either with PBS (for immunolabeling) or with Ames’ solution (for patch clamp recordings). For the experiments on dissociated cells, the hESC-derived RPE monolayers were treated with TrypLE Select for 10 min in +37 °C, gently mechanically triturated with a pipette and centrifuged for 5 min at 1000 rpm. Dissociated cells were resuspended in culture medium, seeded on glass coverslips coated with poly-L-lysine (Sigma-Aldrich) and allowed to settle down for 10 min for patch clamp recordings and 30 min for immunolabeling.

Mouse RPE was prepared for immunolabeling as follows. C57BL/6 mice were euthanized by CO**2** inhalation and cervical dislocation in accordance with the ARVO Statement for the Use of Animals in Ophthalmic and Vision Research and Finland Animal Welfare Act 1986. The eyes were enucleated and bisected along the equator, and the eyecups were sectioned in Ames’ solution buffered with 10 mM HEPES and supplemented with 10 mM NaCl, pH was adjusted to 7.4 with NaOH (Sigma-Aldrich). The retina was gently removed from the eyecup leaving the RPE firmly attached to the eyecup preparation.

### Patch clamp recordings

Ionic currents were recorded from mature hESC-derived RPE monolayers or freshly dissociated cells using the standard patch-clamp technique in whole-cell configuration. Patch pipettes (resistance 5-6 MΩ) were filled with an internal solution containing (in mM) 83 CsCH**3**SO**3**, 25 CsCl, 10 TEA-Cl, 5.5 EGTA, 0.5 CaCl**2**, 4 ATP-Mg, 0.1 GTP-Na, 10 HEPES, and 5 NaCl; pH was adjusted to ~7.2 with CsOH and osmolarity was ~ 290 mOsm (Gonotec, Osmomat 030, Labo Line Oy, Helsinki, Finland). In some experiments, the internal solution also contained 2 mM QX-314-Cl (from Sigma-Aldrich). During all recordings, the tissue was perfused at 2.5 ml min^-1^ with Ames’ solution (Sigma-Aldrich) buffered with 10 mM HEPES and supplemented with 10 mM NaCl and 5 mM TEA-Cl. The pH was adjusted to 7.4 with NaOH and the osmolarity set to ~ 305 mOsm. The bath solution contained 10 nM-10 μM Tetrodotoxin (TTX) citrate (from Tocris Bioscience) when the effect of TTX on the recorded currents was investigated, and 30 μM 18α-glycyrrhetinic acid (from Sigma-Aldrich) when the effect of gap junctional coupling was tested. For the channel subtype recordings, the bath solution was supplemented with 30 nM 4,9-AnhydroTTX, 1 μM A-803467, or 600 nM μ-Conotoxin GIIB. All recordings were made in voltage-clamp mode with pClamp 10.2 software using the Axopatch 200B patch-clamp amplifier connected to an acquisition computer via AD/DA Digidata 1440 (Molecular devices, USA). The access resistance was below 30 MΩ and the membrane resistance above 150 MΩ. Series resistance was 15-30 MΩ and was not compensated. Holding potentials were corrected for a 3 mV liquid junction potential during the data analysis. All recordings were performed at room temperature.

### Immunolabeling

Prior to immunolabeling, samples were washed three times with PBS and fixed for 15 min with 4% paraformaldehyde (pH 7.4; Sigma-Aldrich). After repeated washes with PBS, samples were permeabilized by incubating in 0.1% Triton X-100 in PBS (Sigma-Aldrich) for 15 min and subsequently blocked with 3% BSA (BSA; Sigma-Aldrich) for 1 h. All immunolabeling incubations were done at room temperature.

Primary antibodies against the following proteins were used in this study: cellular retinaldehyde-binding protein (CRALBP) 1:400 (ab15051, Abcam), Connexin 43 (Cx43) 1:200 (C6219, Sigma-Aldrich), N-Cadherin 1:200 (C2542), Nay1.1 1:200 (ASC-001, Alomone labs), Nay1.2 1:200 (ab99044, Abcam), Nay1.3 1:200 (ASC-004, Alomone labs), Nay1.4 1:200 (ASC-020, Alomone labs), Nay1.5 1:200 (AGP-008, Alomone labs), Nay1.6 1:200 (ab65166, Abcam), Nay1.7 1:200 (ASC-008, Alomone labs), Nay1.8 (AGP-029, Alomone labs), Nay1.9 1:200 (AGP-030, Alomone labs), Pan Na_v_ (Na_v_) 1:200 (ASC-003, Alomone labs), Rab7 1:50 ab50533, Abcam), Zonula occludens-1 (ZO-1) 1:50 (33–9100, Life Technologies). All primary antibodies were diluted in 3% BSA in PBS and incubated for 1 h.

The incubation with primary antibodies was followed by three PBS washes and 1 h incubation with secondary antibodies; goat anti-rabbit Alexa Fluor 568, donkey anti-rabbit Alexa Fluor 488, donkey anti-mouse Alexa Fluor 568, donkey anti-mouse Alexa Fluor 488, goat anti-guinea pig Alexa Fluor 568, goat anti-mouse Alexa Fluor 488, donkey anti-rabbit Alexa 647, donkey anti-mouse Alexa 647, goat anti-guinea pig Alexa Fluor 647 and goat anti mouse Alexa Fluor 405 (all from Molecular Probes, Thermo Fisher Scientific) diluted 1:200 in 3% BSA in PBS. Actin was visualized using either a direct phalloidin Alexa Fluor 647 conjugate 1:50 (A22287, Thermo Fisher Scientific), Atto-633 1:50 (68825, Sigma-Aldrich) or tetramethylrhodamine B conjugate 1:400 (P1951, Sigma-Aldrich) and the nuclei were stained with 4’, 6’-diamidino-2-phenylidole (DAPI) included in the mounting medium (P36935, Thermo Fisher Scientific). When primary antibodies from the same host were co-labeled, one was directly conjugated to fluorophore 488 using the APEX conjugation kit (A10468, Thermo Fisher Scientific).

### Phagocytosis assay for hESC-derived and mouse RPE

The porcine POS particles were isolated and purified as previously described^52,53^. Briefly, the eyecups obtained from a slaughterhouse were opened and retinas were removed using forceps under dim red light. The retinas were shaken gently in 0.73 M sucrose phosphate buffer and separated after filtering in sucrose gradient using an ultracentrifuge (Optima ultracentrifuge, Beckman Coulter, Inc., Brea, CA) at 112,400 x g for 1 h at 4°C. The collected POS layer was centrifuged 3000 x g for 10 min, +4°C and stored in 73 mM sucrose phosphate buffer at -80°C.

The purified POS particles were fed to the hESC-derived cells in a KO-DMEM medium supplemented with 10% fetal bovine serum (FBS) and incubated for 2 h at 37° in 5 % CO^2^. In the blockers experiments, selective blockers for Na_v_1.4, Na_v_1.6 and Na_v_1.8 and TTX were also added to the medium for the incubation. Then the monolayers were washed twice briefly with PBS and fixed with PFA according to the immunostaining protocol. Phagocytosis was studied *in vivo* by preparing the mouse eyes under dim red light either at light onset or 15 min, 2 h and 10 h after it. The mice were reared in normal 12-hour light/dark cycle. When blockers were used, the eyecup was opened and then incubated in blocker solutions diluted in Ames’ as described above, for 1 h at 37° with the retina left intact.

### Quantification of bound POS particles in hESC-derived RPE

To detect and quantify bound POS particles, large random fields were imaged with Zeiss LSM780 LSCM on inverted Zeiss Cell Observer microscope (Zeiss, Jena, Germany) by using Plan-Apochromat 63x/1.4 oil immersion objective with 2.0 zoom. The images were first blurred with a Gaussian function after which a Z-maximum intensity projection was binarized using a global threshold. The number of POS particles was then analyzed from the images converted to mask.

### Statistical analysis of the POS phagocytosis quantification

Phagocytosis was repeated three times and the images were pooled together. The normality of the data was tested by using Shapiro–Wilk normality test. Finally, pair-wise comparison was conducted by using two-sided Student’s T-test to confirm the possible statistical significance between the experimental conditions.

### Confocal microscopy and image processing

Confocal microscopy was performed with Zeiss LSM780 LSCM on inverted Zeiss Cell Observer microscope (Zeiss, Jena, Germany) by using Plan-Apochromat 63x/1.4 oil immersion objective. Voxel size was set to x=y=66nm and z=200nm and 1024x1024 pixel stacks of 70-120 slices were acquired with line average of 2. The Alexa Fluor 405 was excited with 405 nm diode laser; Alexa Fluor 488 with 488nm laserline from Argon laser; Alexa Fluor 568 and TRITC with 561nm DPSS or 562nm *InTune* laser; Atto 633 and Alexa Fluor 647 with 633 nm HeNe and 628nm *InTune* laser. Emission was detected with windows of (in nm): 410 – 495 (DAPI, Alexa Fluor 405), 499 – 579 (Alexa Fluor 488), 579 – 642 (Alexa Fluor 568) and 642 – 755 (Alexa Fluor 647). Laser powers were minimized to avoid bleaching and photomultiplier tube sensitivities were adjusted to obtain optimal signal-to-noise ratio of the signal. The data was saved in .czi format and deconvolved using Huygens Essential (SVI, Hilversum, Netherlands) software. The deconvolution was performed with theoretical PSF, signal-to-noise ratio of 5 and quality threshold of 0.01. Information regarding the refractive index of the mounting media was provided by the manufacturer (Vector Laboratories, Burlingame, USA). Images were further processed with ImageJ^54^ and only linear brightness and contrast adjustments were performed for the pixel intensities. Final figures were assembled using Adobe Photoshop CS6 (Adobe Systems, San Jose, USA).

## Acknowledgements

We would like to acknowledge the following contributors. We are grateful to Dr. Jari Hyttinen (Tampere University of Technology) for resources and support. We thank Drs. Kristian Donner (University of Helsinki) and Joshua Singer (University of Maryland) for valuable comments on the manuscript. We acknowledge Outi Heikkilä (Tampere University of Technology), Outi Melin and Hanna Pekkanen (University of Tampere) for technical assistance and Hannele Uusitalo-Järvinen (University of Tampere) as well as Petri Ala-Laurila Lab (University of Helsinki) for providing the animal tissue. University of Tampere Imaging Facility is gratefully acknowledged. This work was supported by the Academy of Finland Grants 260375 (S.N.), 287287 (S.N.), 294054 (S.N.), 267471 (T.O.I.), by the Emil Aaltonen Foundation and by Päivikki and Sakari Sohlberg Foundation.

## Contributions

Conception and design of the study as well as data acquisition, analysis and interpretation was performed by J.K.J, T.O.I., and S.N. The expertise on human embryonic stem cells and RPE differentiation was provided by H.S. The article was written and revised by J.K.J, T.O.I., H.S., and S.N.

## Materials and correspondence

Correspondence to Soile Nymark

